# Amplitude effects allow short jetlags and large seasonal phase shifts in minimal clock models

**DOI:** 10.1101/825331

**Authors:** Bharath Ananthasubramaniam, Christoph Schmal, Hanspeter Herzel

## Abstract

Mathematical models of varying complexity have helped shed light on different aspects of circadian clock function. In this work, we question whether minimal clock models (Goodwin models) are sufficient to reproduce essential phenotypes of the clock: a small phase response curve (PRC), fast jetlag and seasonal phase shifts. Instead of building a single best model, we take an approach where we study the properties of a set of models satisfying certain constraints; here a one-hour pulse PRC with a range of three hours and clock periods between 22h and 26h. Surprisingly, almost all these randomly parameterized models showed a four hour change in phase of entrainment between long and short days and jetlag durations of three to seven days in advance and delay. Moreover, intrinsic clock period influenced jetlag duration and entrainment amplitude and phase. Fast jetlag was realized in this model by means of an interesting amplitude effect: the association between clock amplitude and clock period termed ‘twist’. This twist allows amplitude changes to speed up and slow down clocks enabling faster shifts. These findings were robust to the addition of positive feedback to the model. In summary, the known design principles of rhythm generation – negative feedback, long delay and switch-like inhibition (we review these in detail) – are sufficient to reproduce the essential clock phenotypes. Furthermore, amplitudes play a role in determining clock properties and must be always considered, although they are difficult to measure.

## 1. Introduction

Mathematical modeling in chronobiology has a long tradition [1–5]. In the last few decades, the underlying gene-regulatory networks were identified [6, 7] and subsequently used to design detailed kinetic models [8–10]. Mathematical models helped study entrainment [11, 12], temperature compensation [13], mutant phenotypes [14, 15], jetlag [16, 17], and seasonality [18, 19]. Detailed kinetic models reach enormous complexity if multiple genes and feedbacks are included [15, 20–22].

In this paper, we step back and address the question whether or not simple models can reproduce phenotypic features, such as phase response curves (PRCs), entrainment, jetlag, and seasonality. We ask – what can we learn from a systematic analysis of generic models about the underlying mechanisms? How oscillator properties, such as period and amplitude, govern phenotypic features including jetlag and seasonality? We find that ensembles of quite basic models reproduce these phenotypic features surprisingly well. Moreover, it turns out that amplitudes have a strong effect on jetlag duration and entrainment phase.

As a starting point, we summarize first the major phenomenological features of circadian clocks: PRCs, jetlag, and seasonality. Then we introduce simple models of delayed negative feedback loops with switch-like inhibition.

### 1.1. Phase Response Curves (PRCs) characterize circadian oscillators

Pulses of light or other *Zeitgebers* can shift the phase of endogenous oscillators allowing entrainment [23]. Advancing and delaying phase shifts are quantified traditionally by PRCs [2, 24, 25]. In mammals, even long-lasting bright light pulses induce phase-shifts of just a few hours [26–29]; we term these “small PRCs”. We exploit these observations to parameterize our simple feedback models: the *Zeitgeber* strength in our models is adjusted to get a three-hour difference between maximal delay and advance in response to one-hour light pulses consistent with mouse data [27].

### 1.2. Short jetlags indicate flexibility of circadian clocks

Mammalian clocks are quite strong oscillators [30] and even 6.7h light pulses with 10000 lux can induce only phase shifts of a few hours [26]. Nevertheless, following shifts of day-night cycles by six hours (a common “jetlag protocol”), it is possible to reset the clock within a few days [31–34]. It has been shown previously [35] that global parameter optimization can reproduce small PRCs and short jetlags. Here, we study jetlag duration systematically in generic models without any prior parameter optimization.

### 1.3. Seasonality implies four hour shifts of entrainment phase

Most organisms adapt their intrinsic clock to seasonal variations of day length. In many cases the “midpoint of activity stays relatively stable” [36, 37]. Frequently, phase markers appear to be coupled to dusk or dawn [38, 39]. A change from an 8:16 light-dark (LD) cycle to a 16:8 LD cycle implies phase shifts of about four hours. For example, midnight locked peaks are four hours after light offset in short (8h) nights, but eight hours after offset in long (16h) nights. If morning and evening peaks follow dusk or dawn, their phases with respect to midnight have to be shifted by four hours between winter (8:16 LD) and summer (16:8 LD). Consequently, a four hour change in entrainment phase allows sufficient seasonal flexibility.

### 1.4. Design of generic feedback models

A delayed negative feedback loop with switch-like inhibition is a common mechanism to produce self-sustained oscillations. In the mammalian circadian clock, there are two feedback loops that feature these ingredients [21] (Figure 1A). First, *Bmal1* transcripts are translated to protein that activates *Rev-ErbA* transcription. *Rev-ErbA* after translation to protein inhibits *Bmal1* transcription, thus completing the loop. Second, *Per2* represses its own transcription after delays due to transcription, translation and nuclear import [40]. These two regulatory loops can be abstracted to the ‘Goodwin’ model (Figure 1B), where variables *X*, *Y* and *Z* are involved in a negative feedback loop. We use the Goodwin model, since it is a generic feedback model and can be easily extended to include other features, such as positive feedback (Figure 1C).

**Figure 1:**
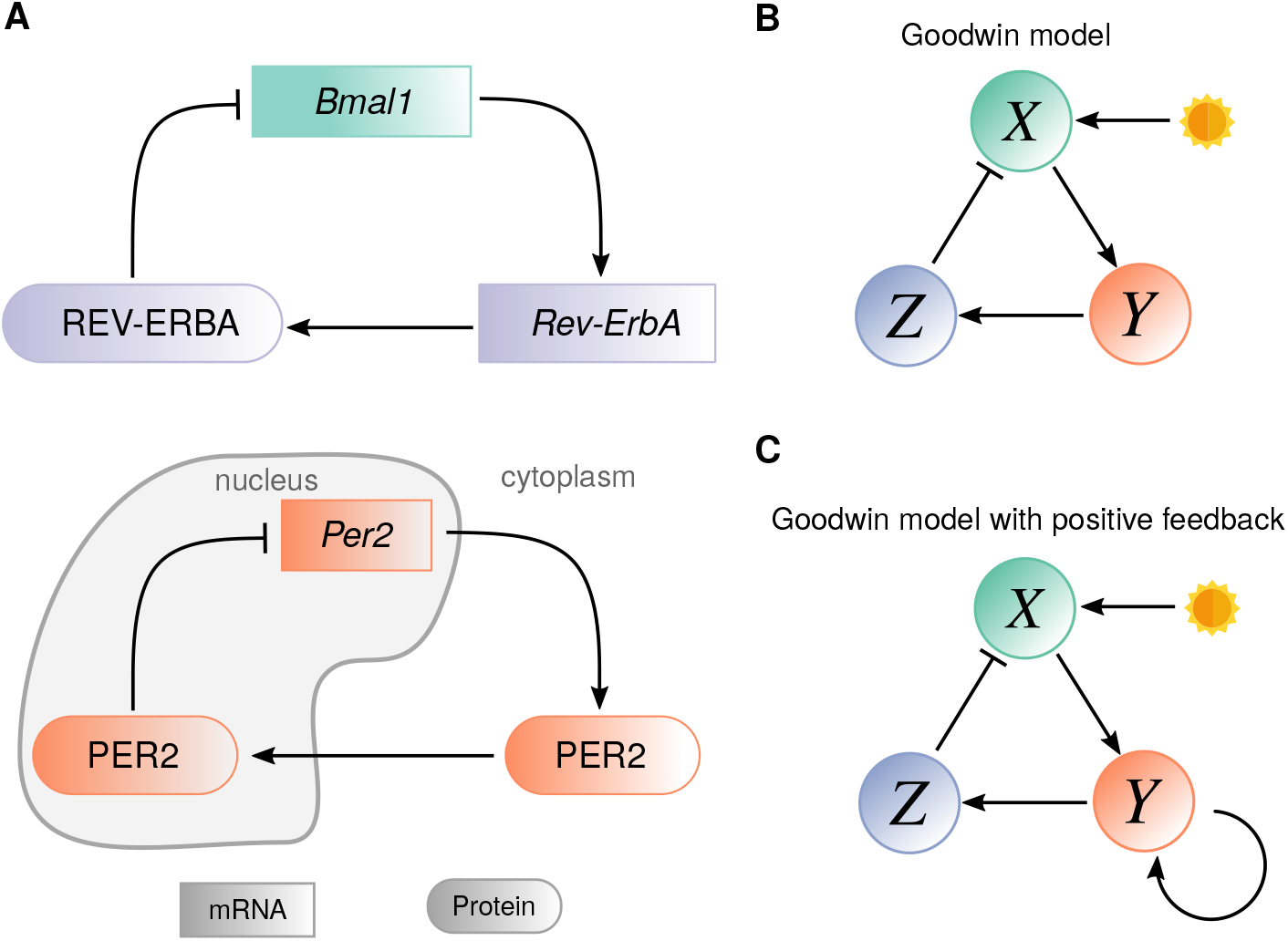
Schematic of minimal models. (A) Two possible gene-regulatory networks in mammalian circadian clock that can produce oscillations via a negative feedback. (B) The three variable model representing the schema in (A) with light input on variable *X* implemented in the simulations. (C) Variant of (B) with a positive feedback on variable Y.

## 2. Results

### 2.1. A representative model reproduces clock features

We explored the behavior of models with the structure in Figure 1B, which is commonly called the ‘Goodwin model’. We varied the six model parameters randomly within a predefined region in parameter space. Several model parameterizations (henceforth shortened to ‘models’) oscillated in a self-sustained manner (Figure 2A is a representative example). The *Zeitgeber* influences the model with an additive term to the production rate of variable *X*. The *Zeitgeber* strength for each model was adjusted to get a three hour range for the phase response curve (PRC) to a one hour *Zeitgeber* pulse (Figure 2B); this served to normalize the entraining input across models. The ‘Material and Methods’ section contains a detailed description of the numerical simulations.

**Figure 2:**
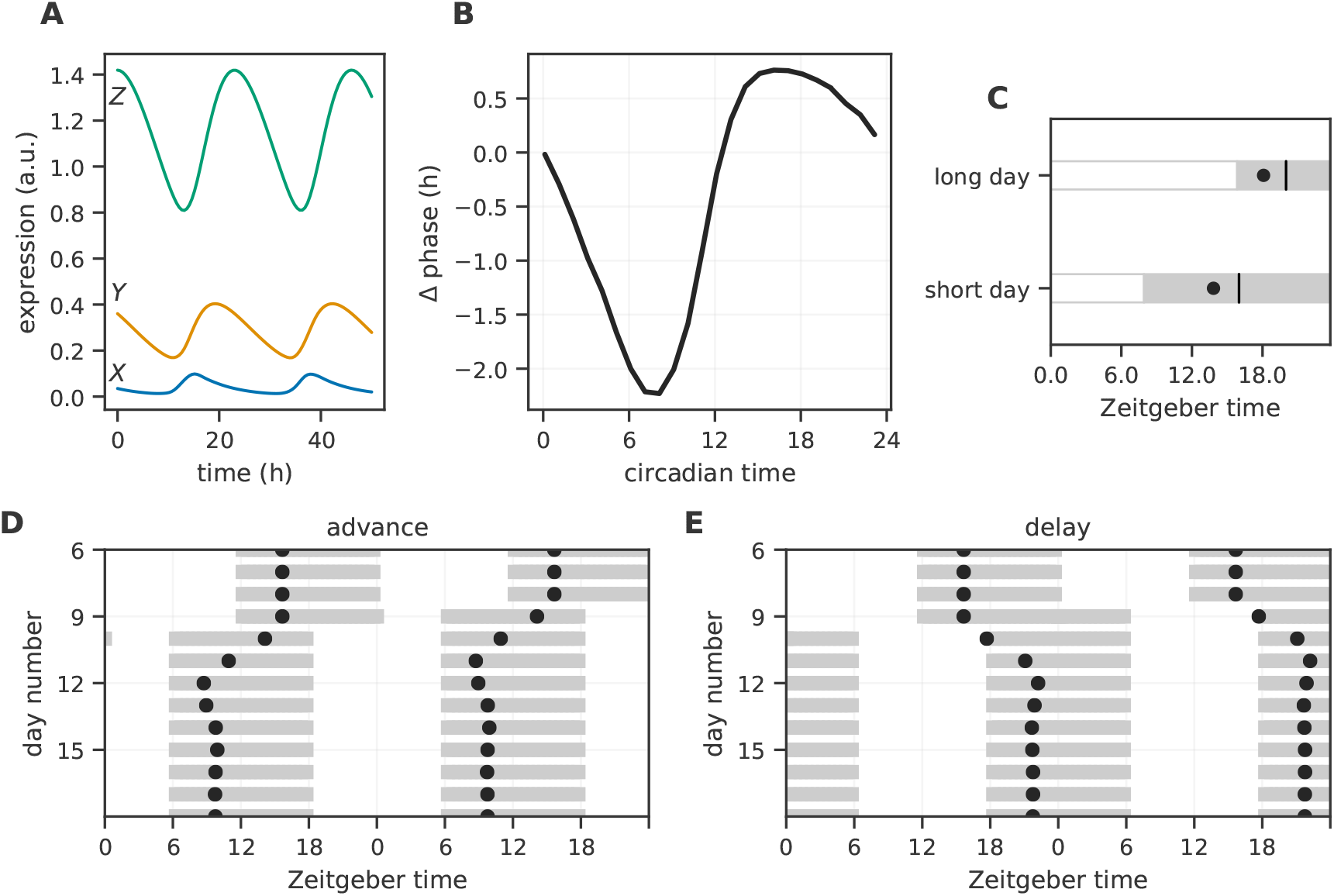
A representative model. (A) Time series of the three variables (marked on the left) of the Goodwin model for a representative parameterization. (B) The phase response curve for the model in (A) in response to a one-hour pulse at different circadian times. (C) The phase of entrainment (black dots) measured relative to lights on under long day (LD 16:8) and short day (LD 8:16). The phase of entrainment is the acrophase of variable *Z*. (D, E) The acrophase of variable *Z* (black dots) during a 6h simulated jetlag from Berlin to New York (delay) or New York to Berlin (advance). Model Parameters: *d*_1_ = 0.15h^−1^, *d*_2_ = 0.15h^−1^, *d*_3_ = 0.25h^−1^, *K* = 0.61, *h* = 11.44, *τ* = 23.03h.

We characterized the seasonal phase shift and duration of jetlag in each model. The seasonal phase shift is the change in the phase of entrainment between short day (LD 8:16) and long day (LD 16:8). The seasonal phase shift of the model in Figure 2A is about four hours (Figure 2C). The phase of entrainment approximately tracks the midpoint of the dark phase.

We also subjected the models to advancing and delaying jetlag of six hours under a 24h (LD 12:12) *Zeitgeber* input (Figure 2D,E). An abrupt change in *Zeitgeber* phase (jetlag) shifts the phase of the model until a stable phase relative to the shifted *Zeitgeber* phase is achieved. The course of this change is the jetlag transient. The jetlag shifts simulated here represent flying from Berlin to New York (delay) or New York to Berlin (advance). The representative model shifted within 4-5 days to within 15 min of the final stable phase for both advancing and delaying jetlags. Moreover, the delaying jetlag shift was shorter than the advancing jetlag shift in this model.

### 2.2. Short jetlag and large seasonal shifts are common

The Monte Carlo approach outlined in the previous section generated an ensemble of models (*N* = 1186). All the models were self-sustained oscillators and had a one-hour pulse PRC with a range of about three hours by design (Figure S1); we allowed a tolerance of 0.25h in order to speed up optimization of the *Zeitgeber* strength. We present here (Figure 3) the landscape of the duration of jetlag and magnitude of seasonal phase shifts within this ensemble.

**Figure 3:**
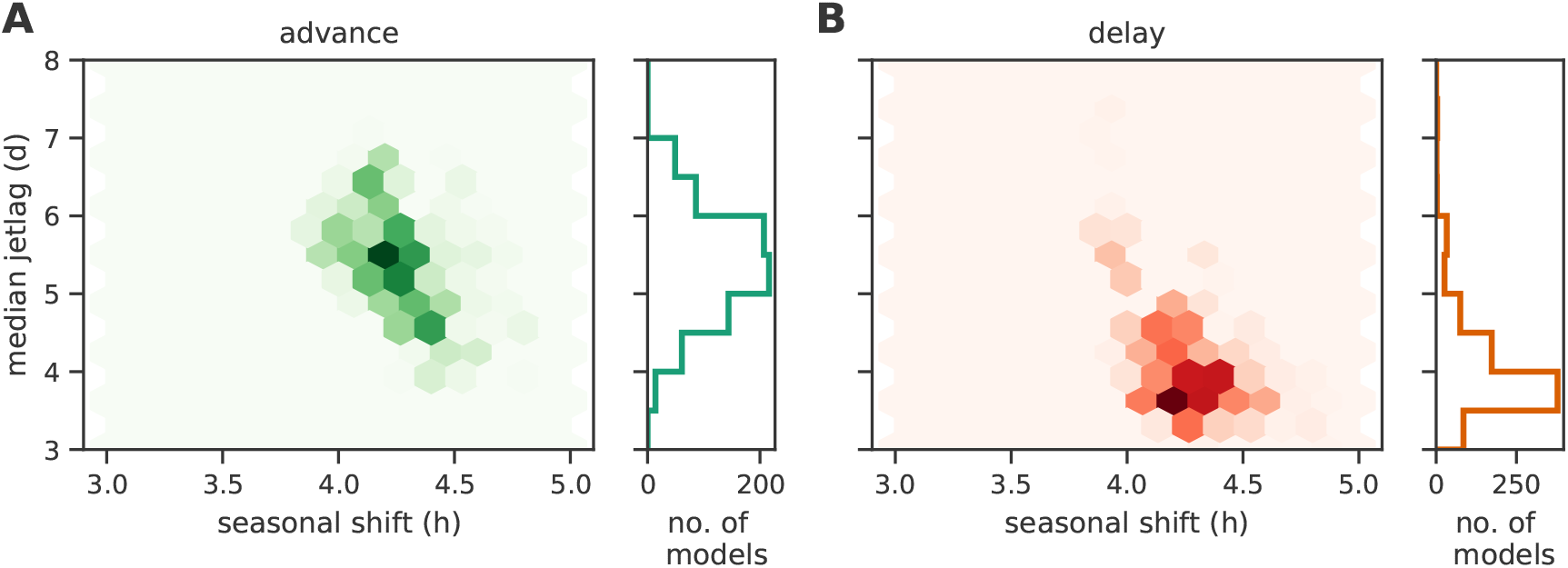
Landscape of jetlag duration and seasonal phase shifts. Combinations of median jetlag for a 6h advance (A) and a 6h delay (B) of the light-dark cycle and seasonal phase shift between long day (LD 16:8) and short day (LD 8:16) observed in the ensemble of models. The number of models for each combination of jetlag and seasonal shift are hexagonally-binned. The counts increase from the lightest to the darkest color.

The duration of the jetlag transient depends on the phase of the *Zeitgeber* at the instant of the jetlag shift in addition to the magnitude and direction of the jetlag shift. Therefore, jetlag was measured as the median jetlag duration starting at four phases six hours apart in order to capture the dependence on the starting phase. Since advancing and delaying jetlags were different (Figure 2), we present results for 6h advancing and 6h delaying jetlags separately.

The whole ensemble exhibited short jetlag shifts (about 3-7 days in length) and seasonal phase shifts of about four hours that matched the observed phenotypes described earlier. We expected specific optimized parameter sets alone to produce the necessary phenotypes [35]. Contrary to our expectation, randomly-chosen parameter sets with the same PRC yielded a highly clustered combination of reasonable jetlag duration and seasonal phase shift.

Advancing jetlag shifts were slower than delaying jetlag shifts (Figure 3). This might be anticipated from the larger delay region compared to an advance region in the PRC (Figure 2B). In fact, one-hour pulse PRCs of all models in the ensemble had approximately a maximum advance of one hour and a maximum delay of two hours (Figure S1). We can roughly estimate (using the PRC) that an advance of one hour or delay of two hours requires a day. If shifts on subsequent days can be carefully adjusted, the simulated 6h jetlags would need six days for the advance or three days for the delay. Surprisingly, the randomly-parameterized models in the ensemble shifted as fast or sometimes faster than the rough estimates. We examine, in the remaining sections, the ensemble properties to better understand the mechanisms that result in the landscape in Figure 3.

### 2.3. Entrainment phases correlate with clock period

The theory of entrainment describes the relationship between the *Zeitgeber* strength, *Zeitgeber* period, intrinsic clock period and intrinsic clock amplitude in a certain clock (model) needed for entrainment [2, 30, 41]. Entrainment has occurred once the clock achieves a fixed phase (called phase of entrainment *ψ*) with respect to the *Zeitgeber*. Instead of observing a single oscillator under a range of *Zeitgeber* periods, we observed the ensemble (comprised of models with different intrinsic clock periods) after entrainment to a 24h (LD 12:12) *Zeitgeber*. The heterogeneity of human circadian clocks was similarly used to study the entrainment properties of humans [42, 43].

Generally, short period clocks had earlier *ψ* and long period clocks had later *ψ* (Figure 4A). The slope of *ψ* versus clock period *τ* was small (slope ~ 0.5) across the ensemble. Unicellular organisms and plants under entrainment show such small slopes [44]. Moreover, entrainment theory predicts ~12h change in the phase of entrainment across the entrainment range of an oscillator [2, 30]. This suggests a large entrainment range is necessary to achieve this small slope. We shall return to the range of entrainment in Figure 8.

**Figure 4:**
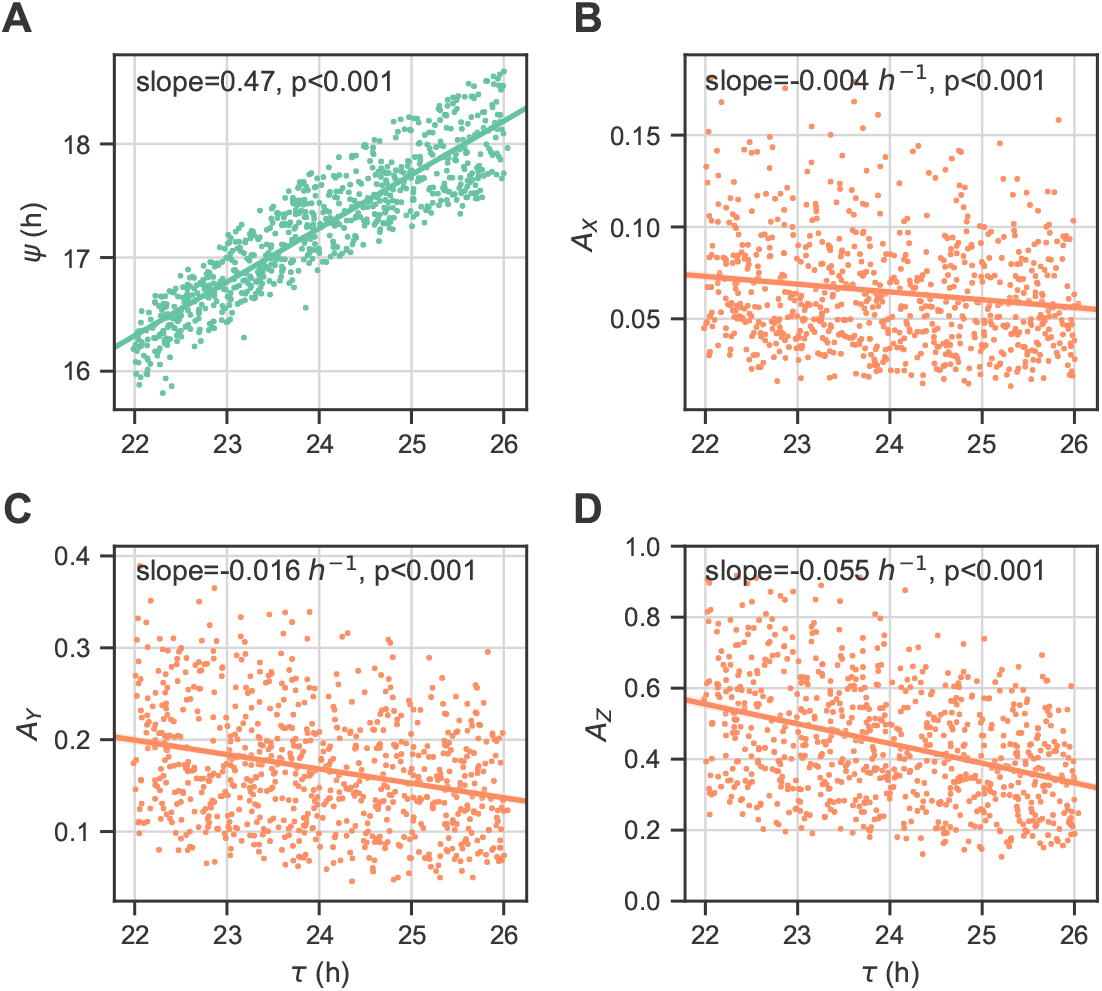
Entrainment properties of the ensemble of models. (A) Dependence of the phase of entrainment *ψ* to a 24h (LD 12:12) *Zeitgeber* on the intrinsic clock period *τ* of models in the ensemble. (B,C,D) Association between the entrained amplitudes of the three model variables *X* (B), *Y* (C), *Z* (D) and the intrinsic clock period of the models in the ensemble. A straight line regressed on the intrinsic period is also plotted.

The entrained amplitudes of the models in the ensemble, on the other hand, showed a small but significant decreasing trend with intrinsic clock period *τ* (Figures 4B-D); such an amplitude trend has been previously reported [45]. Our choice of model parameters (see Materials and Methods) ensured that the intrinsic (unentrained) amplitudes and intrinsic clock periods were uncorrelated in the ensemble.

The observed trend arises from the phenomenon of ‘twist’ in these models; twist is an association between the amplitude and period of an oscillator. The representative model, for example, has a positive association between period and amplitude (Figure 6C). Extrapolating to the whole ensemble, when the *Zeitgeber* period is shorter than the intrinsic clock period, the entrained amplitude decreases (e.g., *τ* = 26h). The corresponding argument for shorter *τ* explains the decreasing trend in data. We will return to this amplitude effect in Figure 6.

### 2.4. Intrinsic periods affect length of jetlag

Both advancing and delaying jetlag were generally short throughout the ensemble. The PRC yielded reasonable rough estimates of the jetlag duration. Using the heterogeneity in the model PRCs within the ensemble, we could test whether the PRC indeed can predict the jetlag duration. If this hypothesis were true, models with larger regions of advance should have shorter jetlag transients to advances and similarly models with larger regions of delay should have shorter jetlag transients to delays. Models whose PRCs have larger delays (and smaller advances, since delay + advance region was chosen to be about three hours) indeed had shorter delaying jetlag transients (Figure S2A). Intriguingly, models with larger regions of delay (and smaller regions of advance) also had shorter advancing jetlag transients (Figure S2B). Therefore, the PRC alone is insufficient to describe the model’s jetlag behavior. To gain further insight into the models, we explored the dependence of the jetlag duration on the model parameter and properties.

Short intrinsic clock periods favor shorter advancing jetlag durations (*ρ*_spearman_ = 0.25, p-value < 0.001, Figure 5A). Similarly, longer intrinsic periods favor shorter delaying jetlag durations, although the effect was less pronounced (*ρ*_spearman_ = −0.10, p-value = 0.008, Figure 5B). During an advancing jetlag transient, the entrained clock needs to speed up (reduce its period/increase its phase velocity) to achieve the new *Zeitgeber* phase. Thus, the shorter intrinsic period (faster clock) naturally aids the jetlag shift. Likewise, response to a delaying jetlag shift requires slowing the clock, which is favored in long intrinsic period clocks.

**Figure 5:**
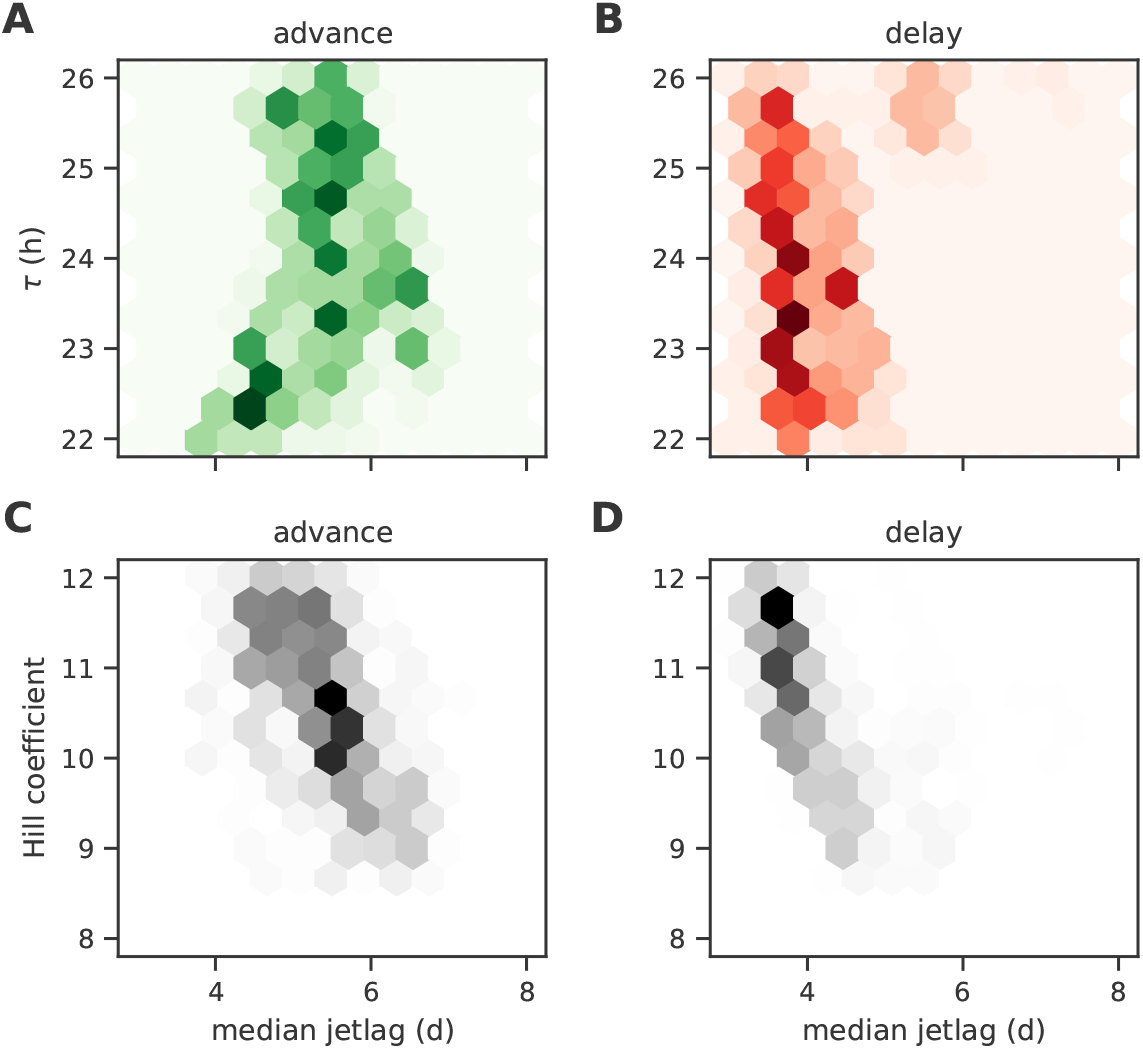
Dependence of jetlag on intrinsic clock period and model nonlinearity. (A, B) Variation of the duration of advance (A) and delay (B) jetlag for models with different intrinsic periods *τ* within the ensemble. (C, D) Influence of the degree of nonlinearity in the feedback loop (Hill coefficient) on the duration of advance (C) and delay (D) jetlags. The number of models for each combination of jetlag and model parameter are hexagonally-binned. The counts increase from the lightest to the darkest color.

**Figure 6:**
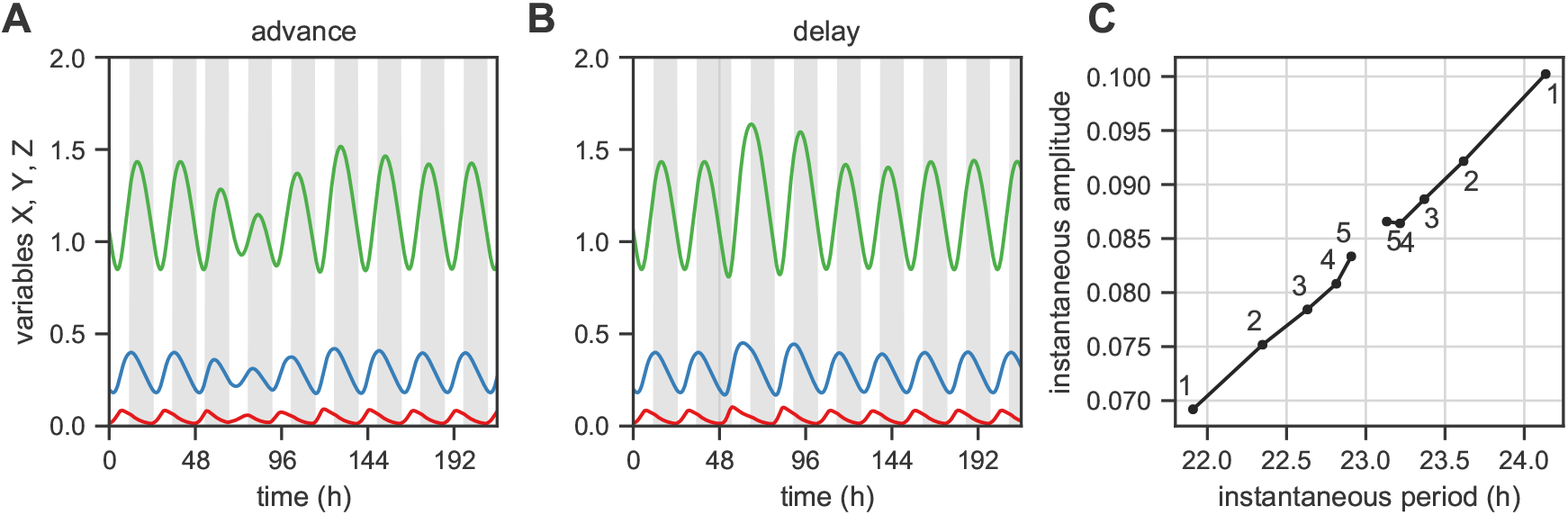
Amplitude changes during jetlag facilitate the phase shift. (A, B) Time series of the representative model in Figure 2 during a 6h jetlag advance (A) and a 6h jetlag delay (B). The dark period of the light-dark cycle is colored grey. Jetlag occurs at *t* = 48h. (C) A measure of the twist (amplitude-period association) of the representative model. The instantaneous amplitude and period of the model as it returns to the unperturbed state after a perturbation of *ϵ* = 0.1 in two opposite directions (the sequence of amplitude and period values is numbered. Only the first five points are shown). Amplitude of variable *X* is used as the amplitude measure.

The Goodwin model requires a Hill coefficient of at least eight in order to oscillate. The Hill coefficient controls the degree of nonlinearity in the negative feedback loop, a necessary component of an oscillator. Models with stronger nonlinearity shifted faster in response to both advancing (*ρ*_spearman_ = −0.56, p-value < 0.001, Figure 5C) and delaying jetlag shifts (*ρ*_spearman_ = −0.75, p-value < 0.001, Figures 5D). The Goodwin model with higher Hill coefficients is associated with higher amplitude relaxation rates and faster entrainment (Figure S3) [46]. Amplitude relaxation rate measures the time taken for the effect of a perturbation to disappear [12, 46]. We suspect that the faster time to entrainment is closely related to the duration of jetlag transient.

### 2.5. Amplitude changes facilitate short jetlag

We earlier observed that both the amplitude and the jetlag duration varied with the intrinsic clock period (Figures 4 and 5). We wondered whether the amplitude of the oscillator played a role in aiding the short jetlags. Therefore, we observed the behavior of the representative model from Figure 2 during simulated advancing and delaying jetlag shifts.

The amplitude of the clock changed significantly over the course of the jetlag transient (Figure 6). The clock amplitude decreased from the steady-state entrained amplitude during a 6h advance (Figure 6A), but returned to the amplitude prior to jetlag after the phase shift had been achieved. On the other hand, the clock amplitudes increased relative to the steady-state amplitudes during a 6h delay. This leads one to question how these amplitude changes aid in shifting the clock phase during jetlag.

To this end, we characterized the *twist* [47, 48] or amplitude-period association of this model without *Zeitgeber* input. We observed how the amplitude and period of the clock varied near the steady-state rhythm (*limit cycle*). The instantaneous amplitude and period of the clock as the perturbed (unentrained) clock returns to the limit cycle is used to measure twist (Figure 6C). It is clear that there is an approximately linear relationship between amplitude and period. Smaller periods are associated with smaller amplitudes and larger periods with larger amplitudes. When the clock phase needs to be advanced, during a jetlag advance, a reduced amplitude is accompanied by a faster clock (i.e., smaller period), which facilitates the advance. Similarly, during a jetlag delay, where phase needs to be delayed, an increased amplitude coincides with a slower clock (i.e., larger period) that aids the delay. This interpretation is consistent with amplitude changes seen in Figures 6A,B.

The amplitude of even the minimal models used here is difficult to define uniquely. The amplitude of each variable in the model can be measured, but it is unclear how these amplitudes can be reduced to a single metric for the whole model. Fortunately, consistent changes in amplitude occur across all three model variables during jetlag and the definition of amplitude does not affect our conclusions.

### 2.6. Observations are robust to the addition of positive feedback

We next investigated the robustness of our observations to changes in the structure of the Goodwin model (Figure 1B). To this end, we added nonlinear degradation of variable *Y* to our base model (Figure 1C) corresponding to proteolysis of the cytoplasmic protein. We previously showed that such a Michaelis-Menten (MM) degradation term implicitly behaves like a positive feedback by reducing the effect of linear degradation [40]. In fact, other processes in the core clock network including complex formation and nuclear import-export are tantamount to positive feedback.

A representative model with positive feedback oscillates (Figure S4A) and has a PRC with about a one hour advance and a two hour delay (Figure S4B), similar to the Goodwin model without positive feedback. The transient after a 6h phase advance or 6h phase delay lasts about four to six days (Figure S4D,E) and the seasonal phase shift between long and short day is also around four hours (Figure S4C).

At the ensemble level, the landscape of jetlag duration and seasonal phase shift (Figure 7A) was tightly clustered like the landscape of the Goodwin model (Figure 3). However, the jetlag duration for both delaying and advancing shifts was slightly longer with positive feedback. Delaying jetlag durations remained shorter than advancing jetlag durations even with positive feedback. The seasonal phase shift was also consistently shorter by about 0.5h with positive feedback. Nevertheless, the qualitative features of short jetlag and seasonal phase shift was robust to these model changes.

**Figure 7:**
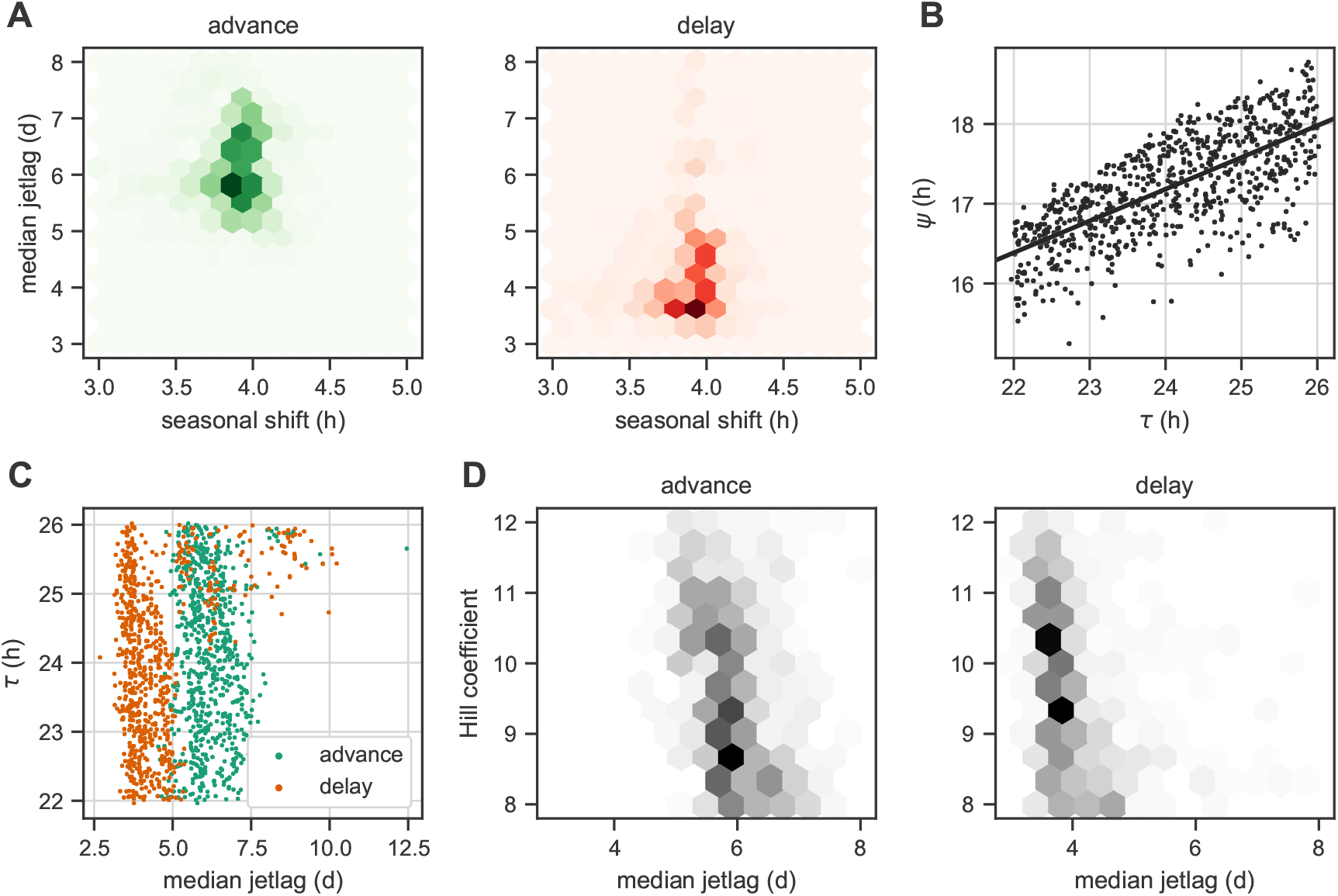
Model phenotypes with an auxiliary positive feedback loop. (A) The landscape of the median jetlag in response to a 6h shift and seasonal phase shift between long day (LD 16:8) and short day (LD 8:16); compare with Figure 3. The advancing (left) and delaying (right) jetlags are shown separately. (B) Dependence of the phase of entrainment *ψ* to a 24h (LD 12:12) *Zeitgeber* on the intrinsic clock period of models in the ensemble; compare with Figure 4A. (C) Variation of the duration of advancing and delaying jetlag (A) for models with different intrinsic periods within the ensemble; compare with Figures 5A,B. (D) Influence of the degree of nonlinearity in the feedback loop (Hill coefficient) on the duration of advancing and delaying jetlags; compare with Figure 5C,D. The number of models for each combination of parameters in (A) and (D) are hexagonally-binned. The counts increase from the lightest to the darkest color.

**Figure 8:**
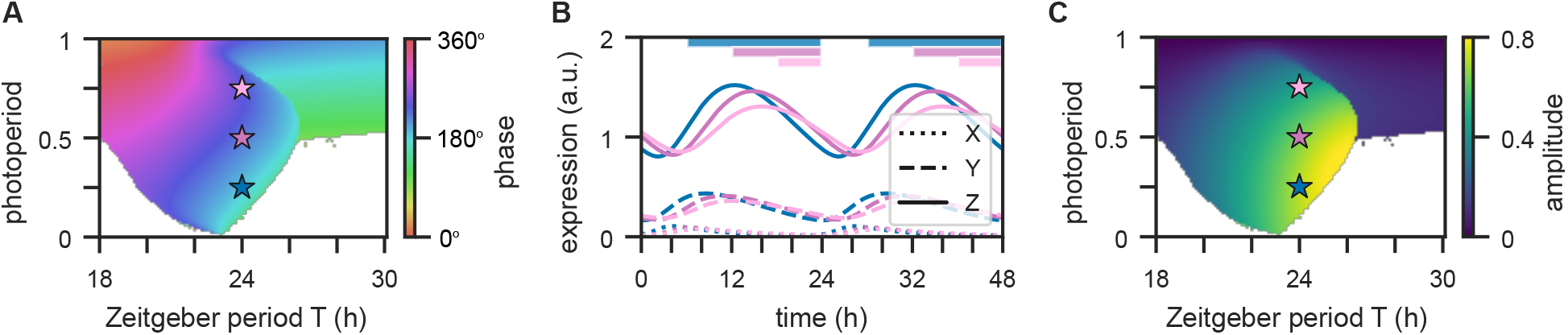
Entrainment properties of the representative model (Figure 2) under varying photoperiods and *Zeitgeber* periods. (A) Entrainment region of the clock model for different *Zeitgeber* periods *T* and photoperiods. Entrainment phases of variable *Z* are color-coded, while regions where no entrainment occurs are depicted by white color. Here, phases (in degrees) are defined as the acrophase of variable *Z* with respect to dawn (lights-on) normalized to the *Zeitgeber* period *T*. (B) Exemplary simulations of the X (dotted lines), Y (dashed lines), Z (bold lines) variables in case of LD 8:16 (blue), LD 12:12 (purple), LD 16:8 (pink) entrainment under a *T* = 24h *Zeitgeber* period. Colored bars denote periods of darkness. (C) Color-coded oscillation amplitudes of the variable *Z* within the entrainment region. *Zeitgeber* period and photoperiod combinations in (B) are with stars of matching color in (A) and (C).

The phase of entrainment *ψ* varied by almost the same slope with positive feedback (Figure 7B). This suggests that the entrainment properties of the model are conserved. When we examined the dependence of short jetlag on the model parameters, the jetlag duration was very weakly affected by intrinsic clock period (Figure 7C). The advancing jetlag transients were unaffected by the circadian period (*ρ*_spearman_ = −0.05, p-value = 0.18), while delaying jetlag transients possessed a weaker reversed trend with respect to intrinsic clock period (*ρ*_spearman_ = 0.11, p-value = 0.002). Higher Hill coefficients consistently favored shorter jetlag duration for both advancing (*ρ*_spearman_ = −0.54, p-value < 0.001) and delaying (*ρ*_spearman_ = −0.52, p-value < 0.001, Figure 7D) shifts. Positive feedback allows oscillations even for smaller Hill coefficients (below eight) [49], which can be seen from the greater spread of Hill coefficients in the ensemble. Furthermore, short jetlag was obtained consistently along a range of Hill coefficients.

Amplitude changes were also observed in the representative model with positive feedback (Figure S4F,G). As with the Goodwin model, jetlag phase advances were accompanied by amplitude reduction and jetlag phase delay by amplitude expansion. This favors short jetlag transients, since the model even with positive feedback has a positive amplitude-period association or *twist* (Figure S4H) [50]. This allows speeding up the clock with amplitude reduction and slowing down the clock with amplitude expansion that can be used to efficiently overcome jetlag.

## 3. Discussion

Circadian rhythms are self-sustained oscillations that can be entrained by light-dark cycles. Mathematical models of these rhythms exhibit limit cycles driven by pulsatile or periodic inputs from a *Zeitgeber*. Any useful model should reproduce known basic phenotypic features, such as the PRC, phase of entrainment and jetlag duration.

In our paper, we studied ensembles of generic feedback models to systematically explore oscillator properties that allow short jetlags and large seasonal phase shifts. These generic models are minimal and contain just six parameters. The *Zeitgeber* strength was adjusted to get a one-hour pulse PRC with a range of three hours [27]. The remaining parameters were randomly sampled within reasonable ranges. Models exhibiting self-sustained oscillations with an intrinsic period between 22h and 26h were analyzed in detail. The resulting set of hundreds of models could be exploited to study connections between oscillator properties and phenotypic features.

Figure 2 demonstrates that a minimal model with a reasonable PRC has large seasonal flexibility and short jetlags. We show in Figure 3 that these features are reproduced by the whole ensemble of models. Furthermore, we analyzed how periods, amplitudes and entrainment phases are related (Figure 4) and studied dependencies of jetlag duration on intrinsic periods and strengths of nonlinearities (Figure 5).

Figure 6 illustrates an interesting mechanism: amplitude changes can affect jetlag duration. It has been described earlier in experimental studies that small amplitude oscillators are easy to reset [12, 51–54]. In these examples mutations and compromised coupling improved resetting and shortened jetlag. In Figure 6C, we point to another amplitude effect: correlations of amplitudes and periods (termed “twist”) imply that *Zeitgeber*-induced amplitude changes can speed up or slow down oscillations. Consequently, transient amplitude changes as shown in Figures 6 and S4 can tune jetlag duration.

In order to evaluate the robustness of our result, we generated another ensemble of models with an auxiliary positive feedback. Figures 7 and S4 demonstrated that parameter dependencies and reproduced features do not change drastically. Moreover, a different auxiliary positive feedback in the form of the saturating degradation of the repressor [55] can produce deadzones often observed in experimental PRCs [56, 57], which could be an interesting next step.

Many of the described results can be quantified by extensive bifurcation analysis of the models [19]. Figure 8 shows an example of a two-dimensional bifurcation diagram for the representative model: the entrainment range, phase (Figure 8A) and amplitude (Figure 8B) are presented for varying *Zeitgeber* periods and photoperiods. It turns out that entrainment phases and amplitudes vary widely. Selected model time courses (Figure 8C) demonstrate again that amplitudes and entrainment phases co-vary.

Another interesting observation is the abrupt widening of the entrainment range for long days. Bifurcation analysis (Figure S5) revealed that this arises from systematic changes due to increased light on long days. As found previously in another model [19, 58], increasing light input can transform self-sustained oscillations to damped oscillations (a reverse “Hopf bifurcation”). Damped oscillations can be driven by any extrinsic periodic forcing and hence the “entrainment range” is huge [59, 60]. However, amplitudes decrease drastically for large *Zeitgeber* periods (Figure 8B).

In the following sections, we discuss general aspects of mathematical models of circadian clocks. We emphasize that models are useful on different levels, that long days and switches are necessary for self-sustained oscillations, and that amplitudes are quite important to understand jetlag duration and seasonal flexibility.

### 3.1. Modeling can be applied on different levels

We analyzed generic models of a negative feedback loop. The required delay was achieved by a chain of reactions and slow degradation rates. The switch-like inhibition was modeled by a high Hill coefficient. Such a Goodwin model was motivated by self-inhibition of the Period and Cryptochrome genes and the *Bmal1*-*Rev-ErbA* loop. However, in contrast to previous models [15, 22, 61, 62], we did not fit models to specific expression data. Therefore our models, while not quantitative in detail, can nevertheless be applicable in a wider context including in organisms other than mammals.

We emphasize that mathematical models of circadian rhythms can be applied on the single cell level describing gene-regulatory feedback loops. Other models might be appropriate on the tissue level, or on the organismic level. The available experimental data are quite heterogeneous. On the genetic level, the phenotypes of many mutants and downregulations are available [63, 64]. On the organ level, many expression profiles have been measured [65, 66]. On the organismic level many data on periods, entrainment phases and PRCs are documented [24, 44]. Consequently, also a great diversity of mathematical models have been developed including phenomenological amplitude-phase models [67], oscillator networks [58], explicit delay models [61, 68] and detailed kinetic gene regulation models [69]. Complex models can reproduce multiple mutants [20, 21] and pharmacological interventions [69].

Here we studied basic Goodwin models [70] that reproduced PRCs, jetlag, and seasonal adaptation surprisingly well. It has been shown earlier that mechanisms of temperature compensation could be also addressed with a simple Goodwin model [13]. Such generic models focus on the most important features of self-sustained oscillations: delayed negative feedback loops and switch-like inhibition.

### 3.2. Circadian rhythm generation requires a 6 hour delay

Under quite general conditions mathematical theory predicts that self-sustained oscillations with a period of 24 hours require a delay of at least 6 hours [61, 71, 72]. Most transcriptional-translational feedback loops introduce delays of about an hour and thus the corresponding periods are in the range of a few hours [73–75]. In order to reach periods of 24 hours, specific extra delays are required. In *Drosophila*, nuclear import of PER and TIM proteins is delayed by about six hours [76]. In mammals and *Neurospora crassa* multiple phosphorylations of the intrinsically disordered proteins PER2 and FRQ contribute to long delays [77–80]. Furthermore, cytoplasmic and nuclear complex formation [81], saturating degradation [82, 83] and epigenetic regulations [84] can induce delays.

### 3.3. Possible mechanisms of switch-like inhibition

In our simple models, a high Hill coefficient is needed to get self-sustained oscillations [49]. Extended models have more realistic exponents due to longer reaction chains [85], positive feedback [40], protein sequestration [86], multi-site phosphorylation [87] and protein inactivation [88]. In any case, switch-like inhibitions are required to get self-sustained rhythms. Theoretical studies have shown that switches can be obtained by cooperativity [89, 90], positive feedbacks [91], competition of antagonistic enzymes [92], and by sequestration [93]. In mammalian clocks, the most essential nonlinearities (”switches”) are not precisely known. It is likely, that multiple phosphorylations [94], complex formations [81], and nuclear translocation [78] play a role. Presumably, the competition of histone acetylation and deacetylation plays an important role since several HATs and HDACs influence clock properties [95]. Moreover, histone modifications are modulated by PER and CRY binding [81]. Similar cellular mechanisms have been described in the *Neurospora* clock [96].

## 4. Conclusions

Our model simulations show that amplitudes have pronounced effects on jetlag duration and entrainment phase. Typically, periods are the main focus of chronobiological studies since there are established devices, such as running wheels or race tubes, to measure periods. The measurement of amplitudes is less straight forward. Quantification of activity records [97] or conidation in race tubes [98] reflects just specific outputs. Gene expression profiles have to be carefully normalized for amplitude quantification [61]. Furthermore it is not immediately evident which gene is representative of the core clock amplitude. In *Neurospora*, FRQ and WCC amplitudes have been considered [99, 100]. In mammals, PER2 [101] and *Rev-ErbA* levels [102] have been used to quantify amplitudes. Measuring the response to pulses provides an indirect quantification of limit cycle amplitudes [101, 103, 104]. Small amplitude limit cycles exhibit larger pulse induced phase shifts than large amplitude limit cycles [25]. Despite the complexities in quantifying amplitude, we suggest studies take into consideration amplitude in addition to the standard phase-based metrics, such as PRCs.

Our comprehensive analysis of generic feedback oscillators provides insight into the design principles of circadian clocks. The essential role of fine-tuned delays and molecular switches is emphasized by mathematical modeling. Reasonable jetlag durations and remarkably large seasonal variations of the entrainment phase can be reproduced by quite simple models. Our models stress the important role of circadian amplitudes. Simulations show that amplitudes contribute significantly to phenotypic features such as jetlag duration and seasonal adaptation.

## 5. Acknowledgements

B. A., C. S. and H. H. acknowledge support from the Deutsche Forschungsgemeinschaft (DFG) (SPP 2041, HE2168/11-1, TRR 186-A16). In addition, C. S. acknowledges support from DFG (SCH3362/2-1).

## 6. Material and Methods

### 6.1. Model structure – Goodwin model

We chose the Goodwin model to represent the core circadian clock feedback loop like many earlier works [13, 58, 105] (Figure 1B). Theoretically, the three variables in this model are the minimum number required to produce self-sustained oscillations. We further reduced the number of parameters in the standard model by using the fact that amplitudes can be rescaled; this is often called nondimensionalization. This yielded a model with five parameters (*d*_1_, *d*_2_, *d*_3_, *K, h*):

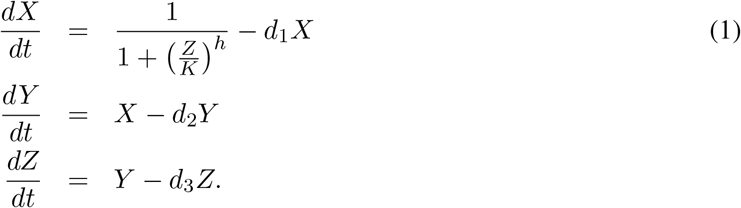

*d*_1_, *d*_2_ and *d*_3_ are degradation parameters with units h^−1^. *h* is the Hill coefficient (unitless) that controls the strength of the nonlinearity in the loop and *K* is the half-maximum concentration of the Hill function controlling the repression of *X* by *Z*.

If the model oscillates for a particular choice of parameters, the period of oscillation *τ*′ is some complex function of the parameters. Scaling the equations produces a model with a desired period of oscillation *τ*:

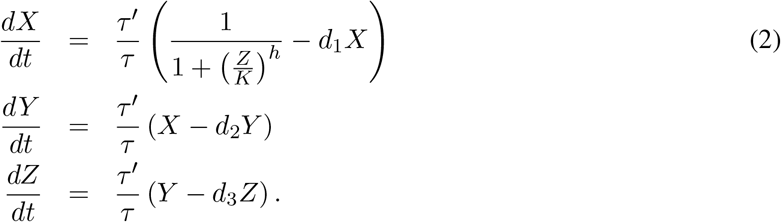

Finally, the input to the clock from the *Zeitgeber* was modeled as an addition to the rate of production of variable *X*. The *Zeitgeber* input consisted of the *Zeitgeber* strength *L* times *I*(*t*), where *I*(*t*) = 1 during the day (light phase) and *I*(*t*) = 0 during the night (dark phase). The model used for studying entrainment, jetlag simulation and seasonal phase shift was:

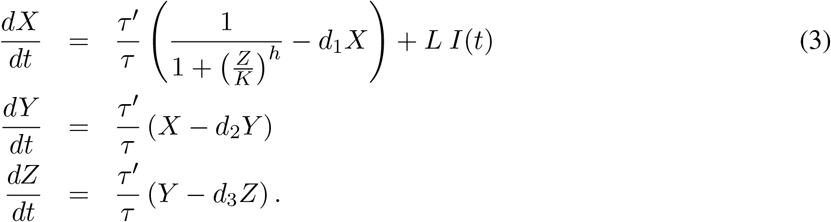

### 6.2. Monte Carlo simulations

In order to study the generic behavior of the Goodwin model, we resorted to the Monte Carlo approach. In this approach, model parameters are chosen randomly to obtain an ensemble of models that can be studied for a variety of phenotypes. We now outline our simulation schemes.

We randomly generated values for the five model parameters. In addition, we also chose a random intrinsic clock period for each model in order to have a uniform distribution of intrinsic periods in the ensemble. In this manner, 2000 different sets of six parameters were generated using latin hypercube sampling (LHS). LHS ensures that parameter sets explore the entire chosen range of each parameter. The degradation parameters could vary between 0.1 and 0.3 to get oscillation periods around 24h. The Hill coefficient could lie between five and twelve. The Goodwin model requires a Hill coefficient of at least eight [49], but the model with positive feedback can also oscillate with smaller Hill coefficients [40]. The half-maximum concentration of the Hill term was chosen between 0.25 and 1 in line with common values taken by the variable *Z* (see (2)). Finally, the intrinsic period was restricted to 22h to 26h based on the range of periods seen in mammalian clocks.

For each random choice of parameter values, we tested whether the model oscillated robustly. If it did, then we optimized the *Zeitgeber* strength *L* such that the one-hour pulse PRC had a range (largest advance-largest delay) of 3 ±0.25h. Bear in mind that scaling the model to reduce parameters makes the oscillator amplitudes arbitrary. The entrainment behavior of the model depends on the ratio between the *Zeitgeber* strength and the oscillator amplitude. We used the constraint on the PRC to keep the effect of the *Zeitgeber* consistent across models in the ensemble.

With the optimized *Zeitgeber* strength *L*^∗^, we ensured that the model can entrain to a 24h (LD 12:12) *Zeitgeber*. Upon entrainment, the phase of entrainment *ψ* was computed as the acrophase of variable *Z* relative to the lights-on (0 to 1 transition) of the *Zeitgeber*. Only if the model oscillates and is entrainable, we computed phenotypes of the model.

For each model, the seasonal phase shift is the difference between the phase of entrainment *ψ* for long day (24h LD 16:8 *Zeitgeber*) and short day (24h LD 8:16 *Zeitgeber*). Furthermore, for each model, we simulated jetlags with both a 6h phase advance and a 6h phase delay. This also ensured that our comparisons were well defined and data for advancing and delaying jetlag shifts are always paired. Since the jetlag duration depends on the *Zeitgeber* phase when the phase shift occurs, we computed the median of the jetlag durations for 6h shifts starting from four different initial *Zeitgeber* phases (0, 6h, 12h, 18h). We measured jetlag duration from the time of the *Zeitgeber* shift to the time point after which the phase of entrainment *ψ* of the clock stayed within 15 min of the final (shifted) phase.

### 6.3. Addition of positive feedback

We have previously defined the basic Goodwin model with different possible auxiliary positive feedback loops [40] and their significance in the mammalian circadian clock. Here, we used a variation of Goodwin model with Michaelis-Menten (MM) degradation of the variable *Y* (Figure 1C):

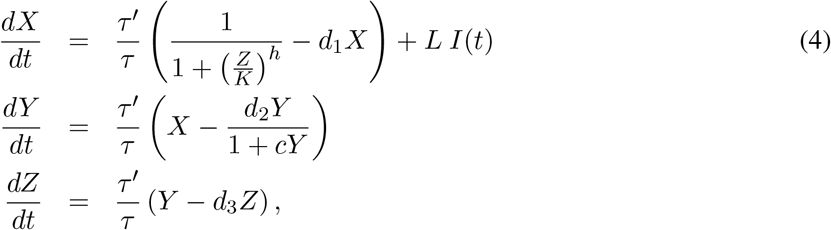

where *c* controls the strength of the positive feedback. In reality, for each simulated standard Goodwin model, we computed the phenotypes for the related positive feedback model (4) by randomly choosing a *c* value between 0.5 and 5. Thus, the comparisons between standard Goodwin and Goodwin with positive feedback models are well-defined, since parameter sets/models are paired. In practice, LHS generated sets of seven variables (five model parameters + intrinsic period + positive feedback strength) and these values were used to simulate the Goodwin model and the corresponding positive feedback model. The computation of model phenotypes was exactly as previously described.

### 6.4. Arnold Onion

The Arnold Onion visualizes the entrainment behavior of the representative Goodwin model (Figure 2) for different combinations of *Zeitgeber* period and photoperiod. For each combination of *Zeitgeber* period *T* and photoperiod, we numerically solved (2) for a total integration time of 2020 entrainment cycles at a step size of Δ*t* = 0.01h. Transient dynamics within the first 2000 cycles were neglected. Next, we saved the complete state of the system at the beginning of the 2001st entrainment cycle and determined whether the system returns to the close neighborhood of this state, given by an *ϵ*-ball of radius 0.01 as described in [19]. Stable recurrence times equal to the *Zeitgeber* period *T* suggests an entrained state of the Goodwin oscillator. This is cross-validated by demanding a small variation of oscillation peak values of the *Z* variable. Phases within the entrained region were determined as the distance between dawn, i.e., lights-on, and the peak time of the oscillations of the Z variable. Amplitudes are determined as half the difference between the peak and trough values.

### 6.5. Floquet Multipiers

The speed at which the model returns to the limit cycle after a perturbation is measured by the amplitude relaxation rate. The average relaxation rate over one cycle can be calculated using the continuation software AUTO07p [106]. The software computes the so-called Floquet multipliers for any model defined using ordinary differential equations. These multipliers then are related to the amplitude relaxation rate.

### 6.6. Bifurcation Analysis

The bifurcation analysis of the model in Figure 2 was performed using XPPAUT with a constant *Zeitgeber* input, whose intensity was varied (Figure S5). Our approach is outlined in [107].

