## Supplemental Figures for "Amplitude effects allow short jetlags and large seasonal phase shifts in minimal clock models"

### Supplementary Figures

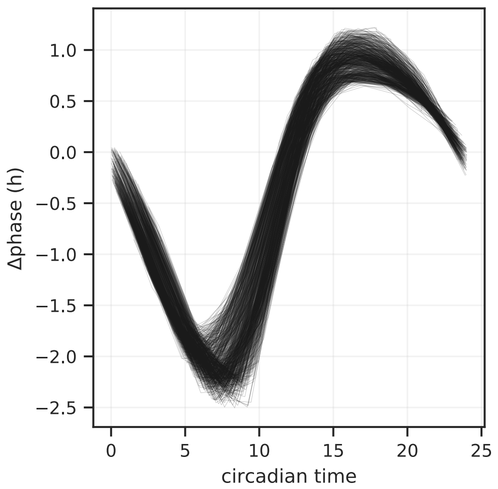


Figure S1: Phase response curves (PRCs) for the set of models. The one-hour pulse PRC for all models in the ensemble with the pulse time scaled by the intrinsic clock period in order to make the PRCs comparable.

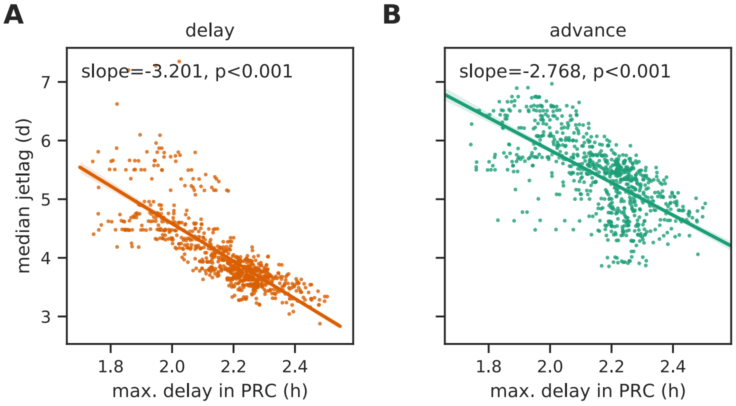


Figure S2: Jetlag duration for models with different PRCs. (A) The dependence of duration of 6h delaying jetlag on the features of the one-hour pulse PRC, i.e., the maximum delay shifts in the PRC. (B) The dependence of the duration of a 6h advancing jetlag on the maximum delay shifts in the PRC. The dependence on the maximum advance of (A) and (B) is just the mirror image, since by design the sum of maximum advance and maximum delay is approximately three hours. The p-value for the slope obtained from linear regression is shown for all plots.

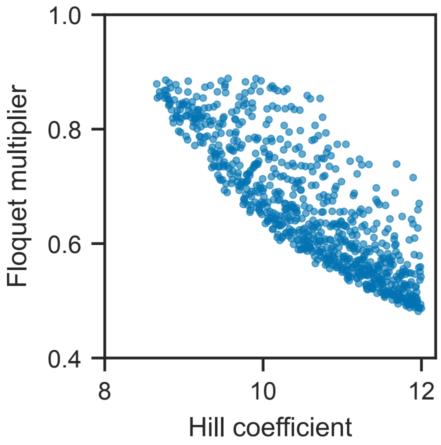


Figure S3: Floquet multiplier of the Goodwin model for different Hill coefficients. The amplitude relaxation rate is $-\log(\text{Floquet multiplier})$.

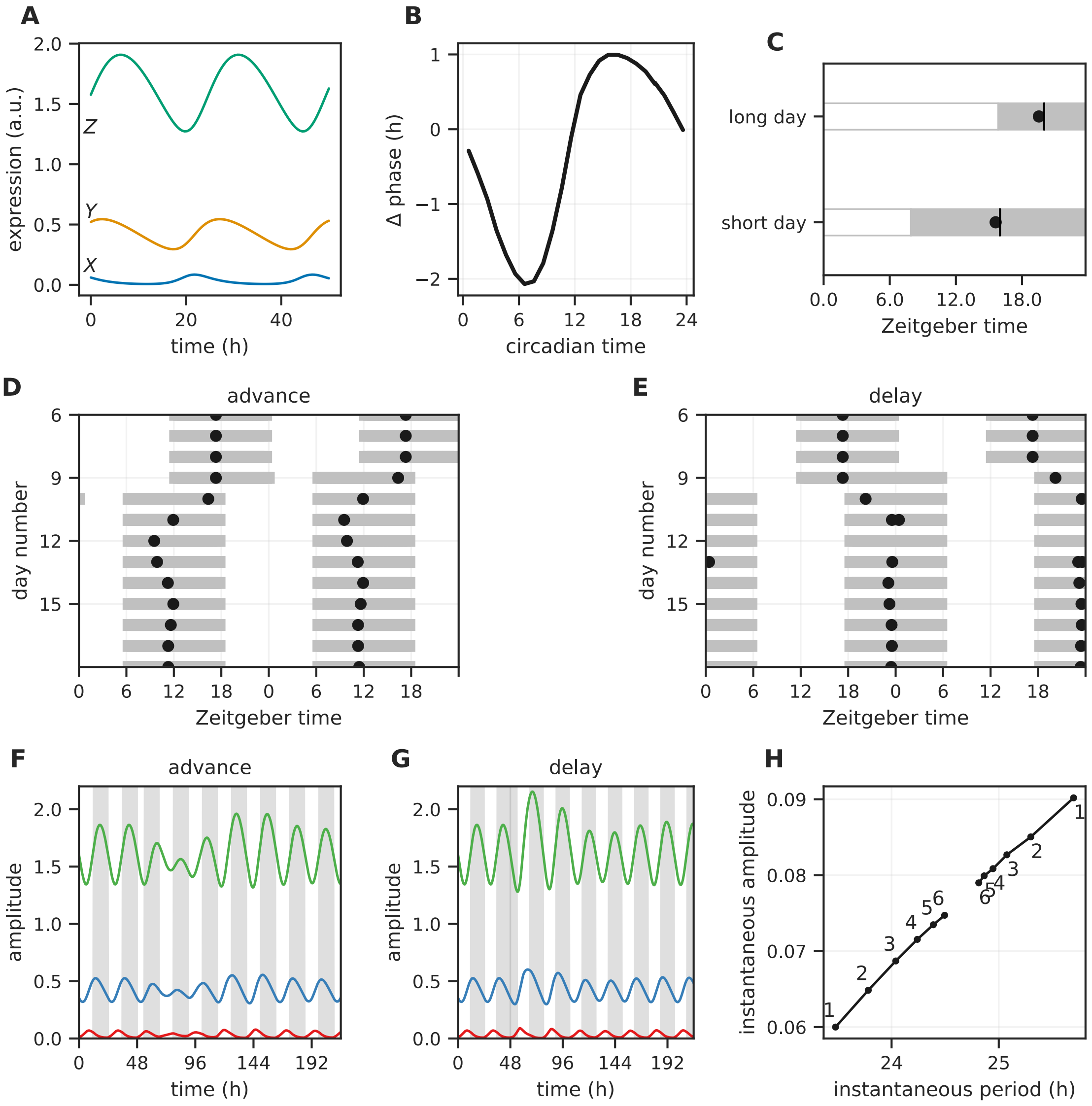


Figure S4: Representative model with positive feedback in the ensemble (compare with Figures 2 and 6). (A) Time series of the three variables of a representative model with positive feedback (Figure 1C). The time series corresponding to each model variable is marked. (B) The phase response curve for the model in (A) in response to a one-hour pulse. (C) The phase of entrainment (black dots) measured relative to lights on under long day (16h light, 8h dark) and short day (8h light, 16h dark). The phase of entrainment is acrophase of variable $Z$. (D, E) The acrophase of variable $Z$ (black dot) during a 6h simulated jetlag from Berlin to New York (delay) or New York to Berlin (advance). (F,G) Time series of the model during a 6h jetlag advance (F) and a 6h jetlag delay (G). The dark period of the light-dark cycle is colored grey. Jetlag occurs at $t \sim 49$h. (H) A measure of the twist (amplitude-period association) of the representative model. The instantaneous amplitude and period of the model as it returns to the unperturbed state after a perturbation of $\epsilon = 0.1$ in a random direction. Amplitude of variable $X$ is used as the amplitude measure. Model Parameters: $d_1 = 0.29\text{h}^{-1}, d_2 = 0.20\text{h}^{-1}, d_3 = 0.26\text{h}^{-1}, K = 0.92, h = 10.01, \tau = 24.71\text{h}$.

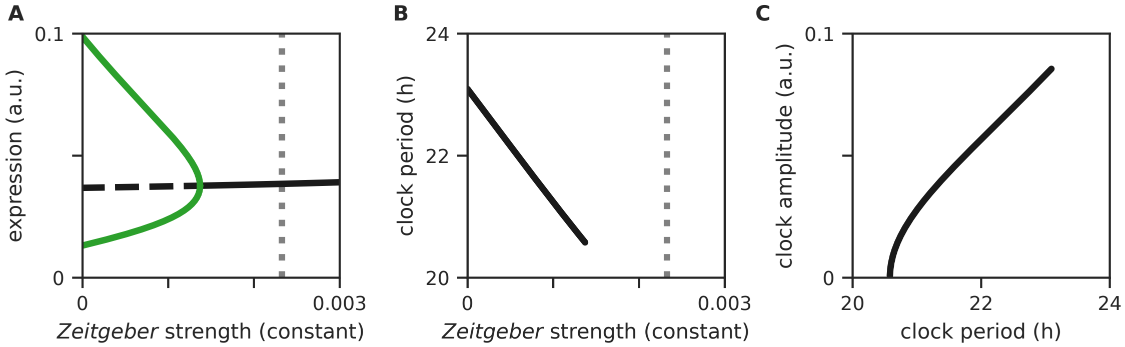


Figure S5: Bifurcation analysis of the representative model from Figure 2. (A) The behavior of the model for different levels of constant *Zeitgeber* input. The stable steady-states are denoted by solid black lines and unstable steady-states by dashed black lines. The green line represents the maximum and minimum expression levels of the $Z$ variable in the limit cycle regime. To the right of this region, only damped oscillations ensue. (B) The period of the model within the limit cycle regime in (A). (C) The amplitude-period covariation of the model within the limit cycle regime in (A). The grey dotted line is the *Zeitgeber* strength used for this model in Figure 8.
